# Resting Parasympathetic Nervous System Activity is Associated with Greater Antiviral Gene Expression

**DOI:** 10.1101/2021.08.21.457209

**Authors:** Danny Rahal, Sarah M. Tashjian, Maira Karan, Naomi Eisenberger, Adriana Galván, Andrew J. Fuligni, Steve W. Cole

**Affiliations:** Department of Psychology, University of California, Los Angeles, Los Angeles, CA 90095, USA; Brain Research Institute, University of California, Los Angeles, Los Angeles, CA 90095, USA; Cousins Center for Psychoneuroimmunology, Semel Institute for Neuroscience and Human Behavior, University of California, Los Angeles, Los Angeles, CA 90095, USA; Department of Psychiatry and Biobehavioral Sciences, University of California, Los Angeles, Los Angeles, CA 90095, USA

**Author notes:** Equal author contribution (Danny Rahal, Sarah M. Tashjian). Correspondence concerning this article should be addressed to Danny Rahal, Franz Hall, University of California, Los Angeles, Los Angeles, CA 90024.

**Keywords:** antiviral gene expression, Type I interferon (IFN) expression, parasympathetic nervous system, autonomic nervous system

## Abstract

Parasympathetic nervous system activity can downregulate inflammation, but it remains unclear how parasympathetic nervous system activity relates to antiviral activity. The present study examined associations between parasympathetic nervous system activity and cellular antiviral gene regulation in 90 adolescents (*M_age_* = 16.3, *SD* = 0.7; 51.1% female) who provided blood samples and measures of cardiac respiratory sinus arrhythmia (RSA), twice, five weeks apart. Using a multilevel analytic framework, we found that higher RSA (an indicator of higher parasympathetic nervous system activity)—both at rest and during paced breathing—was associated with higher expression of Type I interferon (IFN) response genes in circulating leukocytes, even after adjusting for demographic and biological covariates. RSA was not associated with a parallel measure of inflammatory gene expression. These results identify a previously unrecognized immunoregulatory aspect of autonomic nervous system function and highlight a potential biological pathway by which parasympathetic nervous system activity may relate to health.

---

The two branches of the autonomic nervous system—the sympathetic nervous system (SNS) and parasympathetic nervous system (PNS)—reciprocally influence many physiological systems, including the immune system (Kohm & Sanders, 1999; Madden et al., 1994; Nance & Sanders, 2007). Dysregulation of the autonomic nervous system increases risk for inflammatory diseases (e.g., arthritis, athereosclerosis; Carney et al., 2001; Tan et al., 1993; Tsuji et al., 1996), and a substantial body of research has found that the sympathetic nervous system can up-regulate inflammation (Heidt et al., 2014; Powell et al., 2013) and inhibit antiviral biology, particularly expression of the Type I interferons (IFN) that mediate innate antiviral responses (Collado-Hidalgo et al., 2006; Sloan et al., 2007). The genomic impact of this sympathetic immunoregulatory program has come to be known as the Conserved Transcriptional Response to Adversity (CTRA; Cole, 2019). The parasympathetic division of the autonomic nervous system exerts a reciprocal inhibitory effect on inflammatory gene expression (Tracey, 2002, 2009). Less is known about PNS regulation of antiviral gene expression, although there are theoretical grounds to expect that parasympathetic activity might potentially stimulate antiviral biology.

Among its other roles, the autonomic nervous system can serve as a stress response system that mobilizes physiological processes in response to environmental stimuli and threats. When a threat is perceived, PNS activity decreases, and removal of the vagal break enables SNS activation (Porges, 2007). SNS activation in response to acute threats is adaptive, as it enables organisms to mobilize bodily resources to fight or flee the present threat. In the context of the immune system, it may be adaptive for the SNS to promote inflammation, as opposed to antiviral processes, in order to optimize immune responses to wounding injuries (Cole, 2009). This transient, threat-associated martialing of immune resources away from antiviral biology and toward inflammatory responses has been hypothesized to underlie the CTRA profile in leukocytes. Higher levels of this CTRA functional genomic profile are commonly observed in the context of high levels of stress in animals (Chun et al., 2017; Cole et al., 2012; Heidt et al., 2014; Korytář et al., 2016; Powell et al., 2013; Snyder-Mackler et al., 2016; Tung et al., 2012) and have been associated with chronic stress, loneliness, socioeconomic disadvantage, bereavement, early deprivation, post-traumatic stress disorder, and depression in humans (Boyle et al., 2019; Chiang et al., 2019; Cole, 2019; Cole et al., 2015; Miller et al., 2014). Research in animal models suggests that many of these effects are mediated by β-adrenergic receptor activation in response to the SNS neurotransmitter norepinephrine, which alters leukocyte gene regulation pathways to promote transcription of pro-inflammatory genes while reducing transcription of antiviral genes such as Type I IFNs (Cole, 2019; Collado-Hidalgo et al., 2006). As a consequence, β-adrenergic receptor activation is related to enhanced viral replication (Cole et al., 1999, 1998, 2001). Likewise, prior research has also found that social stress increases SNS activity and through this pathway promotes inflammatory and decreases antiviral gene expression (Leschak & Eisenberger, 2019; Powell et al., 2013; Sloan et al., 2007).

Given that the SNS and PNS exert reciprocal control of many physiological processes, it is not surprising that PNS activity downregulates inflammation. Tracey and colleagues have mapped a cholinergic anti-inflammatory pathway (Tracey, 2002, 2009), and experimental studies suggest that *in vivo* stimulation of PNS activity through this pathway inhibits inflammation in rats (Bernik et al., 2002; Bonaz et al., 2017; Borovikova et al., 2000; Jin et al., 2017; Wang et al., 2003). Higher PNS activity also relates to lower production of macrophages in mice (Rosas-Ballina et al., 2011). Likewise, higher heart rate variability—an indicator of PNS activity in humans—has been associated with lower stimulated production of proinflammatory cytokines (Marsland et al., 2007) and lower levels of systemic inflammation in humans (Cooper et al., 2015; Janszky et al., 2004; Sajadieh et al., 2004). Theoretical analyses of the interaction between social processes and autonomic nervous system function also suggest that PNS activity might up-regulate antiviral capacity (Cole, 2009; Leschak & Eisenberger, 2019; Porges, 2007).

By contrast, little is known about the how PNS activity might impact antiviral biology. In this study we examined variation in basal antiviral gene expression (Type I IFN response gene activity) and inflammatory gene expression as a function of individual differences in PNS tone, as measured by respiratory sinus arrhythmia (RSA). Associations were tested in a sample of adolescents aged 14 to 17 years who were recruited as part of a larger behavioral investigation. Prior research has found that resting RSA stabilizes from childhood to adolescence (Dollar et al., 2020; Hinnant et al., 2018), and lower RSA has been linked with clinical illness including depression to a similar extent in adolescents, emerging adults, and older samples (e.g., Koenig et al., 2016; Rottenberg et al., 2007; Yaptangco et al., 2015). For this study, 90 adolescents provided both paced and resting heart rate at two time points five weeks apart and provided blood samples that were assayed for CTRA. Extrapolating from the reciprocal effects of SNS and PNS activity, we predicted that participants with higher RSA would show greater basal Type I IFN activity (primary hypothesis). Based on previous data linking PNS activity to reduced inflammation (Tracey, 2002, 2009), we also predicted that participants with higher RSA activity would show lower levels of inflammatory gene expression (secondary hypothesis).

## Method

### Participants

The present study included a community sample of 90 adolescents (*M_age_* = 16.28, *SD* = 0.73, range 14-17; 51.1% female). Adolescents were recruited from local high schools in Los Angeles through class presentations and flyers. Adolescents completed a prosocial intervention with three conditions, such that there were 30 adolescents with valid blood data collected per condition. Participants were racially and socioeconomically diverse, as indicated by adolescents’ report of race and parents’ reports of annual family income and highest levels of education (Table 1). No participants reported smoking cigarettes, and a minority of adolescents (7.8%) reported ever drinking alcohol. Participants reported low levels of loneliness using the UCLA Loneliness Scale (Russell et al., 1978) and experiencing few stressful life events over the past year using a 14-item checklist (e.g., death of a close family member, begin new school, change in living conditions; Uy & Galván, 2017), and participants’ reports of loneliness and stressful life events were comparable to those of other ethnically diverse community samples of adolescents (e.g., Majeno et al., 2017; Uy & Galván, 2017; Table 1).

**Table 1.** Frequencies for demographic variables.

| Variable | N | %/M(SD) |
| --- | --- | --- |
| Age | 90 | 16.28 (0.73) |
| Female | 46 | 51.11% |
| Sick (Pre-Intervention) | 14 | 15.56% |
| Sick (Post-Intervention) | 10 | 11.11% |
| Income |  |  |
| \$0-\$10,000 | 2 | 2.22% |
| \$10,001-\$20,000 | 3 | 3.33% |
| \$20,001-\$30,000 | 13 | 14.44% |
| \$30,001-\$40,000 | 8 | 8.89% |
| \$40,001-\$50,000 | 6 | 6.67% |
| \$50,001-\$60,000 | 7 | 7.78% |
| \$60,001-\$70,000 | 3 | 3.33% |
| \$70,001-\$80,000 | 6 | 6.67% |
| \$80,001-\$90,000 | 1 | 1.11% |
| \$100,001-\$150,000 | 19 | 21.11% |
| \$150,001-\$200,000 | 9 | 10.00% |
| \$200,001-\$250,000 | 2 | 2.22% |
| \$250,001+ | 10 | 11.11% |
| Mother's Education |  |  |
| Elementary school only | 3 | 3.33% |
| Elementary and middle school only | 3 | 3.33% |
| Middle school completion | 1 | 1.11% |
| Some high school | 0 | 0.00% |
| High school diploma / GED | 7 | 7.78% |
| Some college | 18 | 20.00% |
| Associates degree (2 year degree) | 7 | 7.78% |
| 4 year college degree | 24 | 26.67% |
| Master's degree | 15 | 16.67% |
| Doctorate degree / Medical degree / Law | 6 | 6.67% |
| Father's Education |  |  |
| Elementary school only | 4 | 4.44% |
| Elementary and middle school only | 3 | 3.33% |
| Middle school completion | 3 | 3.33% |
| High school diploma / GED | 7 | 7.78% |
| Some high school | 9 | 10.00% |
| Some college | 12 | 13.33% |
| Associates degree (2 year degree) | 4 | 4.44% |
| 4 year college degree | 21 | 23.33% |
| Master's degree | 13 | 14.44% |
| Doctorate degree / Medical degree / Law | 8 | 8.89% |
| Race/Ethnicity |  |  |
| Hispanic/Latino | 36 | 40.00% |
| European American | 28 | 31.11% |
| African American | 19 | 21.11% |
| Asian American | 9 | 10.00% |
| Middle Eastern | 8 | 8.89% |
| Native American | 5 | 5.56% |
| Other | 9 | 10.00% |
| Biracial | 21 | 23.33% |
| Body Mass Index | 90 | 23.21 (5.41) |
| Loneliness | 90 | 1.96 (0.62) |
| Major Stressful Life Events | 90 | 2.54 (2.19) |

*Note:* Participants could select multiple racial/ethnic backgrounds, such that the percentages sum to over 100%. The possible range was 1-4 for loneliness and 0-14 for major life events.

### Procedures

Data were collected as part of a larger prosocial behavioral intervention, in which participants were randomly assigned to one of three conditions and received text messages every other day for four weeks asking them to do one of the following: to describe their day, complete a kind act for themselves, or complete a kind act for someone else (see Tashjian, et al., 2020 for full details). Participants then completed reports of their kind acts or of their daily activities at the end of the day, and completed a weekly survey regarding their emotion and well-being. Because the present study does not focus on the effects of this behavioral intervention, experimental condition is included as a covariate in all analyses, with the control condition (i.e., reporting daily activities) as the reference group. This intervention involved two laboratory visits: one visit pre-intervention and a second visit five weeks later, one week post-intervention, both scheduled between 8:00 a.m.-12:00 p.m. At each laboratory visit, participants had their heart rate measured and provided blood samples.

Participants had their heart rate measured using electrocardiogram. These data were collected using a physiological recording system with an alternative lead II configuration, with one sensor on each wrist and a third sensor at the top of the sternum, between and below the clavicles (BIOPAC Systems, Inc., Santa Barbara, CA). Participants watched a nature video for four minutes. They then completed a paced breathing exercise, in which they alternated between breathing in for 5 second and breathing out for 5 seconds. They were asked to practice pacing their breathing for three 10-second cycles, and afterward completed the paced breathing exercise for four minutes.

Heart rate data were converted to inter-beat-intervals, and artifacts were edited using CardioEdit by two research assistants who were certified to reliably edit data in this software (Brain-Body Center, 2007). Editing was minimal, and the two research assistants produced nearly identical values (*r* = .99, *p* < .0001). RSA was calculated across a band of high frequencies (.12 - .40 Hz) using the Porges-Bohrer method with the software CardioBatch (Lewis et al., 2012; Porges, 1985; Porges & Bohrer, 1990). High-frequency heart rate variability is used as a measure of RSA because low-frequency heart rate variability can also be influenced by SNS activity (Task Force of the European Society of Cardiology and the North American Society of Pacing and Electrophysiology, 1996). RSA was estimated over 30-second epochs, natural log-transformed to produce normal distributions, and then averaged across epochs (Riniolo & Porges, 2000). By having participants pace their breathing at roughly one breath per 10 seconds, spectral power for HRV is highest at this low respiratory frequency (Kromenacker et al., 2018) such that we are able to calculate an estimate of RSA from the high-frequency spectral power of heart rate variability which is not confounded with respiratory rate. There were no main effects of intervention condition on RSA post-intervention or on changes in RSA between pre- and post-intervention, *p*’s > .10.

Participants also provided a blood sample collected by a phlebotomist at each laboratory visit. Participants reported whether they felt sick (0 = not sick, 1 = sick). Participants also reported the degree to which they were currently experiencing six symptoms including nasal congestion, muscle aches, upset stomach, hot/cold spells, poor appetite, and coughing/sore throat using a five-point Likert scale (1 = not at all, 2 = slight, 3 = mild, 4 = moderate, 5 = severe), in line with a previous study of cellular gene expression (Nelson-Coffey et al., 2017). At both time points, participants reported low levels of symptoms (*M* = 1.36, *SD* = 0.38, range 1-2.5 at first visit; *M* = 1.29, *SD* = 0.33, range 1-2.3 at second visit). Mean ratings for each individual symptom were consistently low, with at least 84% of participants reporting no or slight levels of each symptom at each time point.

Peripheral blood mononuclear cells (PBMC) were isolated by ficol gradient centrifugation and stored at −70°C for subsequent transcriptome profiling in a single batch. Total RNA was extracted from samples using an automated nucleic acid processing system (QIAcube; Qiagen), checked for suitable mass (> 50 ng by NanoDrop One spectrophotometry; achieved *M* = 5042 ± *SD* 2219 ng) and integrity (RNA integrity number > 3 by Agilent TapeStation capillary electrophoresis; actual *M* = 7.9 ± *SD* 0.6) and assayed by RNA sequencing in the UCLA Neuroscience Genomics Core Laboratory using Lexogen QuantSeq 3’ FWD cDNA library synthesis and multiplex DNA sequencing on an Illumina HiSeq 4000 instrument with single-strand 65-nt sequence reads (all following the manufacturer’s standard protocol). Analyses targeted >10 million sequence reads per sample (achieved mean 10.5 million), each of which was mapped to the RefSeq human genome sequence using the STAR aligner (achieved average 93% mapping rate) to generate transcript counts per million total transcripts (TPM; Dobin et al., 2013). Six samples failed endpoint quality control analyses, and all observations from those 6 individuals were deleted from further analysis. TPM values were normalized based on 11 standard reference genes (Eisenberg & Levanon, 2013), floored at 1 normalized TPM to reduce spurious variability, log2-transformed to reduce heteroscedasticity, and *z*-score standardized within gene.

## Analytic Plan

Mixed effect linear model analyses were conducted relating average expression of a pre-specified set of 34 Type I IFN indicator genes (Fredrickson et al., 2015) to RSA (either resting or paced breathing) while controlling for time point (0 = pre-intervention, 1 = post-intervention; repeated measure), indicator gene (repeated measure), and intervention condition (dummy-coded for three conditions). Primary analyses examined unadjusted associations of antiviral gene expression with each RSA measure separately (collected during resting or paced breathing), and secondary analyses additionally adjusted for age (grand-mean centered), sex (0 = female, 1 = male), self-reported potential illness (0 = not sick, 1 = sick), BMI (grand-mean centered), and race/ethnicity (indicator variables for Black, Hispanic Asian, Native-American, Middle-Eastern, vs. reference group Caucasian). Ancillary adjusted analyses additionally controlled for mRNA markers of major leukocyte subsets to control for the potential heterogenetic in leukocyte subset distributions (*CD3D, CD19, CD4, CD8A, FCGR3A, NCAM1, CD14*). A final set of analyses included both resting and paced breathing RSA to determine whether each form of RSA is independently related to antiviral activity. The secondary hypothesis involving inflammation was assessed by models testing associations between resting or paced RSA with 19 pro-inflammatory genes (Fredrickson et al., 2015) and for broader CTRA genomic profile, which was calculated as the difference between inflammatory and antiviral gene expression (i.e., inflammatory – Type I IFN). Analyses were conducted using SAS v9.3 PROC MIXED, specifying a random subject-specific intercept.

## Results

Characteristics of the sample are given in Table 1. Participants primarily identified as Hispanic or Latino (27.78%), biracial (23.33%), or European American (21.11%), with smaller percentages identifying with other ethnic backgrounds. They were socioeconomically diverse, such that roughly half of participants had parents who completed a 4-year degree (50.01% for mother’s education, 46.66% for father’s education). Participants’ RSA values were highly correlated between pre- and post-intervention (*r*’s = .70 - .74), and resting and paced RSA values were also correlated both pre- and post-intervention (*r*’s = .60; Table 2). A small minority of participants reported feeling sick at either the first study visit (*n* = 14;15.6%) or the second study visit (*n* = 10; 11.1%). Parent study experimental condition was unrelated to antiviral gene expression (Table S1), and experimental condition was controlled in all analyses. Levene’s test of residual variation indicated that there were no individual cases of unusually large linear model residual values (i.e., outliers) for this sample across all models, *t*(5150) = 0.14, *p* = .8913.

**Table 2.** Descriptive statistics and correlations between RSA values.

| Variable | *M* | *SD* | *Min* | *Max* | 1. | 2. | 3. | 4. |
| --- | --- | --- | --- | --- | --- | --- | --- | --- |
| 1. Paced RSA Pre-Intervention | 7.59 | 0.93 | 5.15 | 9.59 | — |  |  |  |
| 2. Paced RSA Post-Intervention | 7.46 | 1.00 | 4.42 | 9.45 | .74 | — |  |  |
| 3. Resting RSA Pre-Intervention | 6.96 | 0.98 | 4.42 | 9.64 | .60 | .57 | — |  |
| 4. Resting RSA Post-Intervention | 6.92 | 1.09 | 3.59 | 9.44 | .44 | .60 | .70 | — |

Note: All $p$ 's < .001. RSA=Respiratory Sinus Arrhythmia.

As hypothesized, higher resting RSA, indicative of greater parasympathetic activity, was associated with greater Type I IFN response gene expression, *B* = 0.09, *SE* = 0.03, *p* = .0004. Likewise, higher paced breathing RSA was associated with greater Type I IFN response gene expression, *B* = 0.14, *SE* = 0.03, *p* < .0001 (Fig. 1). In both cases, associations were maintained after adjusting for demographic and biological covariates (Tables S1-S2). When examining both resting RSA and paced RSA in the unadjusted model simultaneously, paced RSA continued to show a significant association with Type I IFN response gene expression, *B* = 0.12, *SE* = 0.03, *p* = .0004, whereas no significant association emerged for resting RSA, *B* = 0.04, *SE* = 0.03, *p* = .2287 (Table S3). The effect of paced RSA remained significant over and above demographic and biological covariates, *B* = 0.10, *SE* = 0.03, *p* = .0022. Although it was still smaller in magnitude than the association for paced RSA, the effect of resting RSA became significant after controlling for demographic and biological covariates, *B* = 0.07, *SE* = 0.03, *p* = .0245.

**Figure 1.**
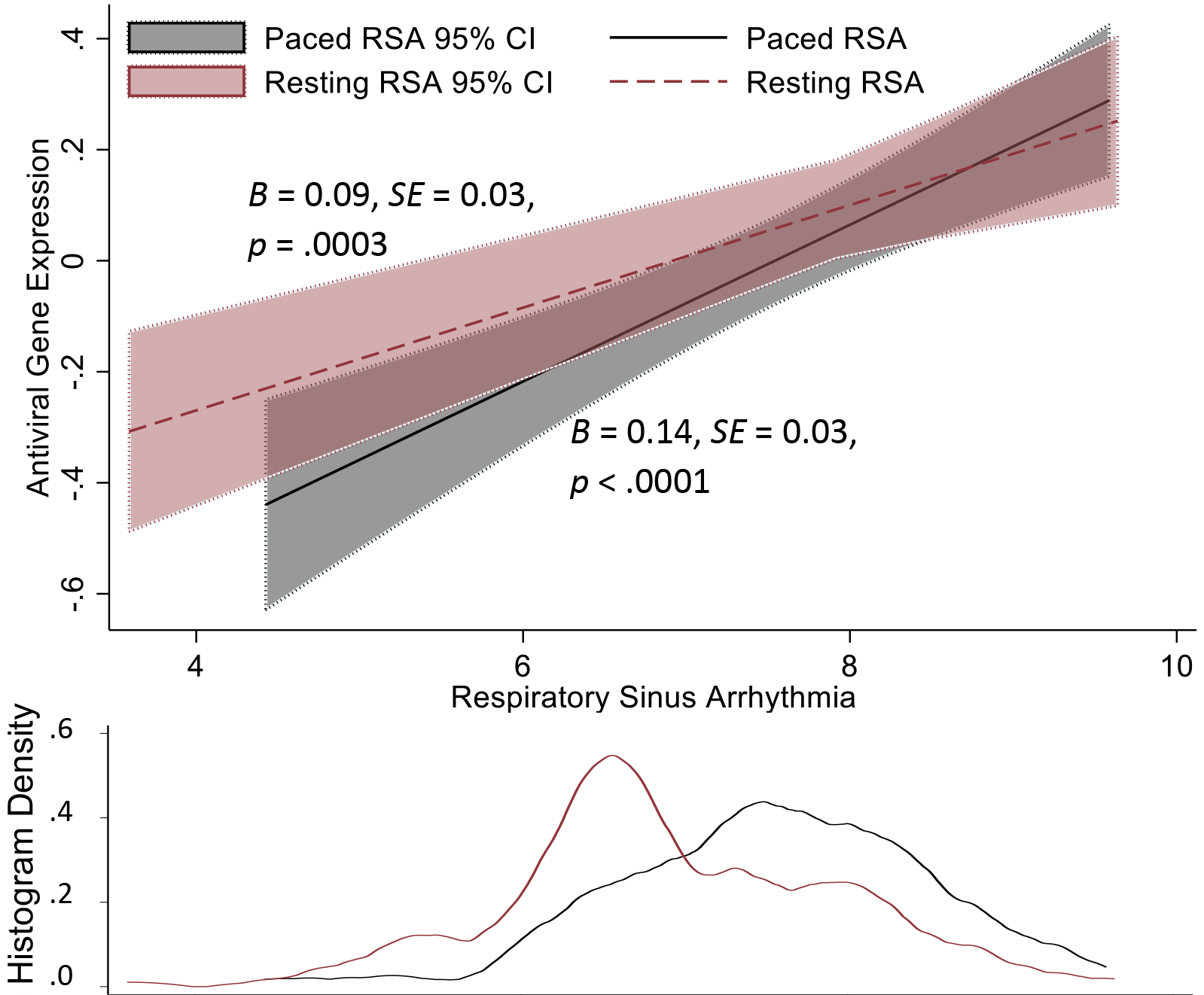
Antiviral gene expression as a function resting and paced RSA. *Note*: Regression lines represent the association between average expression of 34 Type I IFN-related gene transcripts (log2 transformed and z-score standardized) and either resting RSA (dashed line) or RSA during paced breathing (solid line), controlling for time (0 = pre-intervention, 1 = post-intervention) and parent study experimental condition (control condition as reference group). Shaded regions represent 95% confidence intervals for regression lines, which are plotted over the observed range of variation in RSA values for each measure (with RSA distributions shown by color-coded histogram density plot on the lower panel).

Neither resting nor paced RSA showed any significant association with inflammatory gene expression; *B* = 0.03, *SE* = 0.03, *p* = .2364 for paced RSA, and *B* = −0.01, *SE* = 0.03, *p* = .6930 for resting RSA (Fig. S1). Associations remained non-significant after adjusting for demographic and biological covariates (Tables S1-S2).

As would be expected based on the significant differences in Type I IFN gene expression, higher resting and paced RSA were associated with lower expression of the broader CTRA profile (i.e., inflammatory – Type I IFN) for both resting RSA, *B* = −0.06, *SE* = 0.02, *p* = .0022, and paced RSA, *B* = −0.05, *SE* = 0.02, *p* = .0188. These associations remained significant in adjusted models (Tables S1-S2).

## Discussion

Results of the present study show higher PNS activity (as indexed by heart rate RSA) to be robustly associated with greater expression of Type I IFN response genes and, in turn, lower CTRA gene expression profiles in adolescents. No significant associations were observed between RSA and inflammatory gene expression. Although future research will be required to confirm the clinical significance of these observations (e.g., for rates of viral infection and disease), the present results provide preliminary evidence that PNS activity is related to immune regulation, beyond its previously documented effects on inflammatory biology (Tracey, 2002, 2009), and may provide a novel physiological pathway for enhancing host resistance to infectious diseases.

We found that both higher RSA during rest and higher RSA during paced breathing were individually associated with greater Type I IFN-related gene expression. This study is among the first to examine associations between RSA and antiviral biology, particularly among an adolescent sample, and the robust associations across both resting and paced RSA suggest that our findings were not driven by individual differences in respiratory rate (in which case paced RSA would not show associations). Given that SNS and PNS activation often have opposing effects on physiology, our findings also align with previous evidence that greater SNS activity is related to lower antiviral gene expression (Cole et al., 1998, 1999, 2001). The PNS and SNS respond to threatening environmental cues in concert to optimize physiological flexibility (McEwen, 2007), and further mechanistic research is needed to determine whether the PNS directly influences antiviral activity, independent of the SNS (e.g., via parasympathetic neurotransmitter activity), or acts indirectly by antagonizing SNS effects on gene expression. Future research is also needed to compare the strength of associations between antiviral gene expression and resting SNS and PNS activity, and identify strategies to promote optimal antiviral defenses by jointly impacting PNS and SNS activity.

Interestingly, RSA was not related to inflammatory gene expression in this analysis. This finding was at odds with prior literature linking higher PNS activity to lower inflammation through the cholinergic anti-inflammatory pathway (Tracey, 2002, 2009). However, whereas previous research on the cholinergic anti-inflammatory pathway typically examined stimulated immune responses (e.g., by lipopolysaccharide), this study specifically examined baseline RSA rather than RSA in the context of stress or immunologic stimulation. Although our findings suggest that baseline RSA is not related to inflammatory gene expression, acute changes in PNS activity can modulate inflammation, in line with experimental studies (e.g., Bonaz et al., 2017; Jin et al., 2017) and as observed in pre-adolescents (Alen et al., 2021). It is possible that PNS activity is related to inflammation downstream of gene expression. This study was conducted in a sample of adolescents, and adolescents tend to have lower levels of inflammation as compared to adults (De Ferranti et al., 2006). Therefore, it is also possible that associations between PNS activity and inflammatory gene expression may emerge later in development. It is also possible RSA is in fact associated with reduced levels of basal inflammatory gene expression, but such effects were not sufficiently large to reach statistical significance in a sample of this size.

Given the association of RSA with higher expression of Type I IFN response genes, it is not surprising that RSA was also associated with lower CTRA transcriptome profile as antiviral gene expression is the primary inverse component of the CTRA profile (Cole, 2019). Higher levels of CTRA genetic profiles have been linked with stress and poorer mental health among adolescents (Chiang et al., 2019) and adults (Boyle et al., 2019; Cole, 2019; Cole et al., 2015; Miller et al., 2014). These findings align with prior work suggesting that lower RSA is a risk factor for poorer mental health throughout childhood and adulthood and mortality in adulthood (Graziano & Derefinko, 2013). It is possible that lower RSA may predict poorer health through its influence on immune control of viral gene expression. For instance, individuals with HIV tend to show lower PNS activity (McIntosh, 2016), which may contribute to poorer prognosis through reduced antiviral immune response.

Associations between higher RSA and higher antiviral gene expression were robust for both resting and paced RSA when examined separately. However, in simultaneous analyses paced RSA remained the only significant correlate of Type I IFN gene expression. Although RSA is generally considered a valid measure of PNS activity (Beauchaine, 2015; Porges, 1986), there is some discrepancy regarding the interpretation of resting RSA and the need for controlling for respiratory rate or imposing a paced breathing paradigm (Denver et al., 2007; Song & Lehrer, 2003). The consistency of our results across both forms of RSA suggests that associations were robust to variations in RSA assessment, but analyses indicated a stronger quantitative relation for paced RSA. This is consistent with previous observations that paced breathing improves reliability of indicators of PNS activity (Pinna et al., 2007). Therefore, although associations are present between resting RSA and antiviral gene expression, future studies of antiviral gene expression may be best positioned to identify associations by accounting for respiration rate and variability.

Although the present data do not involve any direct measures of clinical health, one potential implication of these findings is that increasing RSA (e.g., through biofeedback or physical training; Sherlin et al., 2009; Wong & Figueroa, 2021) might potentially improve adolescents’ antiviral activity. Lower RSA has also been linked with psychopathology and poorer cardiovascular health (e.g., Beauchaine et al., 2019; Gangel et al., 2017), and the immunological associations observed here might potentially contribute to such relationships. These data also raise the question of whether alterations in PNS activity might potentially be involved in mediating the effects of behavioral interventions that have already been found to reduce CTRA gene expression (Nelson-Coffey et al., 2017), and/or psychosocial risk factors associated with CTRA (e.g., Cole, 2019; Cole et al., 2015).

The current study had key strengths including two time points, an ethnically diverse sample, and assessment of RSA during both rest and paced breathing. However, findings are limited by aspects of the design. The most significant limitation is the fact that causal conclusions cannot be drawn from this observational study relating RSA to antiviral gene expression. Future research experimentally manipulating RSA will be required to document causal effects. The health significance of the observed differences in antiviral gene expression also remain to be documented in future research. Participants completed a behavioral intervention which may have influenced results, although this seems unlikely given that we used both pre- and post-intervention data, the intervention conditions had no detectable effect on gene activity, and results were maintained when controlling for time and intervention condition. Additionally, the sample was limited to adolescents, who are generally low in inflammatory and antiviral gene expression compared to adults (e.g., De Ferranti et al., 2006). It is possible that associations may differ in populations with greater immune activity or in people who have or are at high risk for obesity. Findings should be further assessed among diverse youth, such as those with financial hardship and high levels of adversity who may be at heightened risk for physiological dysregulation with respect to the ANS and immune system (e.g., Johnson et al., 2017; Sloan et al., 2005). Although this study involved an ethnically diverse sample, associations between RSA and antiviral gene expression remained largely unchanged and statistically significant in analyses that controlled for race/ethnicity. Nevertheless, it remains possible that ancestry-related genetic differences (e.g., expression quantitative trait loci) might contribute to the observed associations between higher RSA and greater antiviral gene expression, and future research should integrate measures of genetic polymorphism and ancestry that were not available in this study. Finally, the study lacks a measure of SNS activity. Therefore, we cannot know whether PNS activity relates to antiviral activity uniquely from SNS activity, or whether these systems relate to antiviral activity through similar mechanisms.

Taken together, we found that greater PNS activity—as indexed by higher RSA at rest and during paced breathing—is related to greater antiviral gene expression in adolescents. Greater PNS activity was also related to lower expression of the CTRA, an immune cell transcriptome profile associated with stress and poor mental health, and this association was driven predominately by the antiviral component, with no significant association between PNS activity and the inflammatory component. Results highlight a novel pathway by which youth with lower PNS activity—potentially related to stress and disadvantaged environments—may be positioned for poorer physical health, particularly in an era of increased exposure to viral pathogens.

